# Therapeutic Potential of Dichapetalin M in Metastatic and ER-positive Breast Cancer: Evidence from Cell Line Studies

**DOI:** 10.64898/2026.05.19.724853

**Authors:** George Yankson, Kezia Yaa Awortwe, Mary Anti Chama, Lily Paemka

## Abstract

**Background:** Dichapetalin M (Dic M), an active compound extracted from medicinal plants in the *Dichapetalum* genus, has been previously shown to possess anti-proliferative activity against cancer cell lines. However, the specific mechanism through which it exerts its anticancer effects remains unknown.

**Purpose:** This study focused on elucidating the mechanism of action of dichapetalin M to further explore its potential as a therapeutic agent for resistant and metastatic breast cancer.

**Method:** We confirmed the Estrogen Receptor (ER) as a target of Dic M, using an *in vitro* approach. Furthermore, we examined both the apoptotic and migrastatic effects of dichapetalin M by assessing its impact on the expression of key apoptosis-related and cancer cell migration genes. Finally, we evaluated the compound’s effect on Multi-drug Resistance Gene *MDR1* expression, a gene linked to cancer drug resistance.

**Results:** Our target validation experiments demonstrated that Dic M exhibited considerably higher cytotoxicity in ER-positive breast cell lines compared to ER-negative cell lines. Furthermore, treatment of MCF-7 cells (which are ER-positive) with Dic M led to a dose-dependent increase in the expression of *AREG (amphiregulin)*, a downstream effector of the Estrogen Receptor. Additionally, Dic M inhibited actin polymerization and significantly downregulated genes involved in the turnover of actin monomers. Scratch-wound assay results further demonstrate that Dic M reduces the rate of cell migration, although its impact on EMT-related gene expression was only observed at high doses. Additionally, Dic M treatment in MCF-7 cells resulted in a significant decrease in the expression of pro-apoptotic genes and *MDR1* expression.

**Conclusions:** These findings indicate that Dic M likely interacts with the Estrogen Receptor and employs the apoptotic pathway to exert its cytotoxic and anti-proliferative effects. Dic M exhibits promising potential, such as anti-migrastatic properties and downregulation of a key breast cancer resistance gene, warranting further investigation.

## Introduction

Breast cancer is the second most commonly diagnosed cancer worldwide, posing a significant global health challenge (Bray *et al*., 2024). Its molecular heterogeneity has led to a classification system based on hormone receptor expression, including the estrogen receptor (ER), progesterone receptor (PR), androgen receptor (AR), and the human epidermal growth factor receptor (HER) (Shaath *et al*., 2021). Among these, ERα-positive (ERα+) breast cancers are the most prevalent subtype (Lin *et al*., 2020). Estrogen plays a crucial role in ER alpha-positive (ERα+) breast tumors by acting as a key regulator. When estrogen binds to its receptor, ERα+, it influences the expression of downstream effector molecules, such as cyclin D1, which promote cell survival and proliferation (Clusan *et al*., 2023).

Therapeutic strategies for ER+ breast cancer primarily involve selective estrogen receptor modulators (SERMs), such as tamoxifen, which block estrogen’s interaction with ERα. The application of SERMs is now limited by drug resistance and non-specific side effects (Patel & Bihani, 2018; Chang, 2012). Resistance has been linked to the pregnane X receptor (PXR, also known as NR112), a nuclear receptor overexpressed in cancer tissues. PXR modulates the expression of genes involved in drug metabolism, such as multidrug resistance gene 1 (*MDR1*) and cytochrome P450 3A4 (*CYP3A4*), contributing to reduced therapeutic effectiveness (Chen *et al*., 2007; Skandalaki *et al*., 2021). The activation of PXR is mediated by diverse compounds, and its role in breast cancer drug resistance highlights the need for alternative therapeutic strategies (Abachi *et al*., 2017). In response to these challenges, newer drugs such as fulvestrant (Figure 1) have been developed. Fulvestrant competes with estrogen for ER binding, leveraging its structural similarity to the natural ligand (Nathan & Schmid, 2017). However, emerging evidence suggests breast cancer resistance to fulvestrant (Kaminska *et al*., 2021), underscoring the ongoing need for more effective therapies.

**Figure 1:**
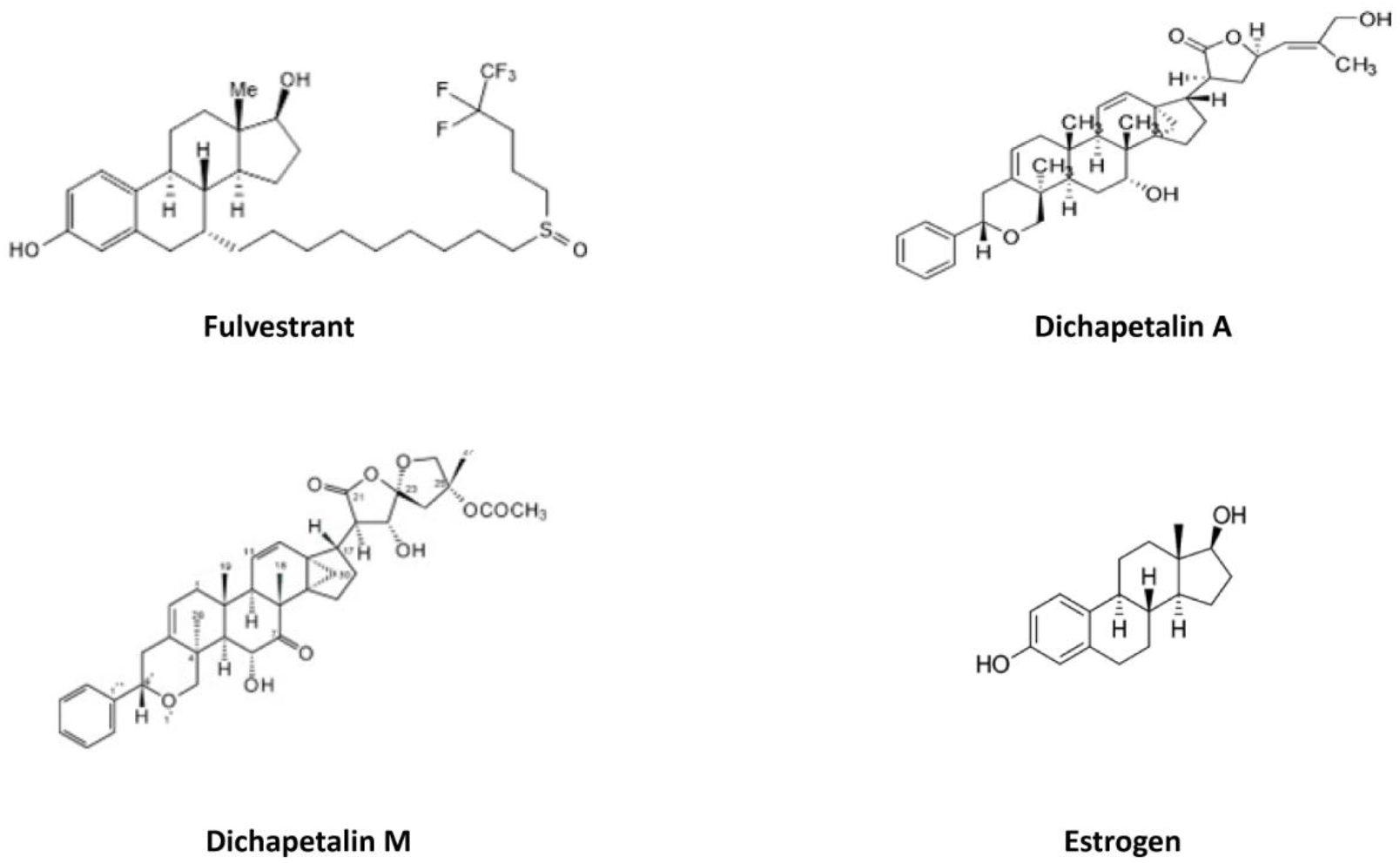
Compounds with triterpenoid moieties similar to those of the estrogen core.

Medicinal plants from the *Dichapetalum* genus offer a promising avenue for drug discovery. Active compounds from plants of this genus, particularly dichapetalins, exhibit a range of biological activities such as anti-tumor and anti-microbial activities (Addae-Mensah *et al*, 2024). Dichapetalins are characterized by a triterpenoid moiety and a 2-phenylpyrano moiety, with structural variations defined by side-chain groups such as lactone, methyl ester, or spiroketal (Osei-Safo *et al*., 2012). Notably, some dichapetalins have demonstrated cytotoxic activity against human cancer cell lines (Fang *et al*., 2006; Osei-Safo *et al*., 2017; Addae-Mensah *et al*, 2024). For instance, dichapetalin A showed activity against various cancer cell lines, including human colorectal carcinoma (HCT116) with EC_50_ = 0.25 μM and towards human melanoma (WM266-4) with EC_50_ = 17 μM (Long *et al*, 2013). Dichapetalin M (Dic M) has demonstrated cytotoxic activities of EC_50_ 0.0099 μM against HCT116 and 0.078 μM against WM266-4 (Long *et al*, 2013). Our previous study (Chama *et al*., 2021) also revealed the activity against MCF-7 by both dichapetalins A (2) and M (3) (Figure 1). *In silico* analysis by Chama et al. (2021) showed that dichapetalins A and M bind to both the ER and PXR receptors. The interaction of dichapetalin M with PXR was further validated in cell-based assays (Chama *et al*., 2021). However, the precise mechanism of dichapetalin M’s anti-proliferative activity against cancer cells remains unknown.

This study is a follow-up on our previous studies on the cytotoxic effect of dichapetalin M (Chama *et al*., 2021), which aims to elucidate the mechanism of action of Dic M in mediating its anti-proliferative effect in breast cancer cell lines. Specifically, we investigated the selectivity of Dic M for ER, assessed its anti-migratory effects, and evaluated its impact on the expression of apoptotic genes and multidrug resistance gene 1 *MDR1*. By bridging these insights, we aim to provide a foundation for the potential development of Dic M as a novel therapeutic agent for breast cancer.

## Materials and Methods

### Cell Lines and Culture

Breast cancer cell lines (MCF 7, MDA-MB 231 and MDA-MB 453) (Table 1) were purchased from ATCC (USA), were cultured and maintained in Dulbecco Modified Eagle Medium (Gibco, USA) supplemented with 10% Fetal Bovine Serum (Sigma-Aldrich, USA) and 1% Penicillin and streptomycin (Gibco, USA) at 37°C in a humidified atmosphere at 5% CO_2_.

**Table 1:** Cell lines and Phenotypes.

| Cell line | PXR | ER |
| --- | --- | --- |
| MCF 7 | + | + |
| MDA-MB 231 | + | - |
| MDA-MB 453 | basal | basal |

(+) receptor expressed (-) receptor not expressed (**basal**) low expression

### Compounds/Drugs

Dic M was isolated from *Dichapetalum madagascariense* at the Department of Chemistry, University of Ghana, as described in our previous studies (Chama *et al*, 2021). Rifampicin, 17β-estradiol (E2), and tamoxifen, which were used as controls, were purchased from Glentham (USA). A metabolite of tamoxifen, 4-OH tamoxifen, which was suitable for *in vitro* studies, was purchased from Sigma-Aldrich (USA) to also serve as a control.

### Determination of the antagonistic effect of Dic M

To determine the antagonistic effect of Dic M on the receptor target, ER, qRT-PCR was conducted to measure the effect of the compound on the expression of *amphiregulin*, a downstream effector molecule of the estrogen receptor. Total RNA was isolated from seeded cells treated with various compounds (E2, 5%, 10%, 25%, and E2+25% Dic M, and vehicle control, 0.01% DMSO) using Zymo Quick RNA extraction kit (Zymogen). The extracted RNA was immediately converted to cDNA using the iScript cDNA synthesis kit (Bio-Rad). Using Luna qPCR master-mix, a total volume of 10μl per well reaction mixture was set up and sealed properly before spinning at 500 rpm for 1 min to remove bubbles. The samples were run on the QuantStudio thermocycler (Applied Biosystems) at cycling conditions recommended by the manufacturer. ACTβ (β*-actin*) served as a reference gene. The various targets were amplified and compared to the corresponding expression in the cells treated with vehicle only (0.01% DMSO). Results from the qPCR were analyzed using the ΔΔC_T_ method. Data obtained were plotted and statistically analyzed using GraphPad Prism software (version 7).

### Determination of the selectivity of Dic M

The selectivity index (SI) of Dic M was calculated by comparing the IC_50_ obtained from its cytotoxic effect on the ER-positive cell line, MCF-7, to that obtained from its effects on ER-negative cell lines (MDA-MB 231 and MDA-MB 453).

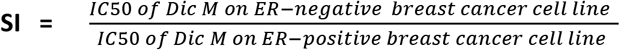

### Effect of Dic M on Actin integrity

MCF-7 cell lines were cultured on a chambered glass coverslip in appropriate media until they were 35-40% confluent at 37°C in a 5% CO_2_ incubator. Cells were treated with varying concentrations of dichapetalin M (5%, 10%, and 25% of the predetermined IC_50_ obtained at 72 hrs of treatment) and incubated for 24 hrs. The complete medium was removed, and cells were washed twice with 1 ml 1X PBS for 5 s. After the washes, cells were fixed with 3.7% paraformaldehyde for 10 mins and permeabilized with 0.5% Triton-X for 5 mins. Phalloidin conjugated with a green fluorescent Alexa Fluor488 dye was prepared by diluting 3.5μl of 14μM of stock in 500μl PBS, and an aliquot of 200μl was added to the permeabilized cells, which were kept in the dark for 30 mins. After washing the cells with 1X PBS, the cells were counterstained with 200μl of 100 nM DAPI. The coverslip was inverted onto a glass slide with 10 μl of mounting media and then sealed with clear nail polish. The slides were observed under a fluorescent microscope (Olympus BX-41) at x100, and images were taken at several XY points. The images obtained were then analyzed using Image J (Schneider *et al*.,2012). Cells treated with vehicle only (0.01% DMSO) were used as negative controls. The effect of Dic M on the expression of factors such as *CFL1* (cofilin 1), *PFN1* (profilin 1), and *LIMK1* (LIM Domain Kinase 1), which are involved in regulating the turnover of actin monomers, was assessed using qRT-PCR.

### Effect of Dic M on cell migration

The scratch-wound assay protocol developed by Mouritzen & Jenssen (2018) was used to investigate the ability of Dic M to inhibit cell migration in a metastatic breast cancer cell line, MDA-MB 231. Cells were seeded at a density of 5.0 × 10^5^ cells/ml for 24 hrs, after which spent media was removed and the wound was created using a 200μl pipette tip and a rule. Cells were gently washed twice with PBS to remove the detached cells. This was followed by treatment with Dic M dissolved in DMEM media supplemented with 1% FBS. Images were taken at ×100 using the Optika microscope at different time points (2 hrs, 4 hrs, 6 hrs, 12 hrs, and 24 hrs). Images taken were then analyzed using the *Wound Healing Size Tool* plugin in ImageJ (NIH, Bethesda, MD, USA) to assess the change in wound area.

### Influence of Dic M on apoptosis

The cytotoxic effect of dichapetalin M was determined by assessing the expression of genes in the intrinsic apoptosis pathway. The expression of the pro-apoptotic genes *BAD* and *BIM*, and the anti-apoptotic gene, *BCL-2*, following Dic M or vehicle only treatment (0.01% DMSO) was investigated using RT-qPCR as described above. With *ACTB* (β-actin) as the reference target, the results obtained were analyzed using the ΔΔCT method. The sequences of the primers used are listed in Supplementary Table S1.

### Assessment of drug resistance modulation by Dic M in breast cancer cell lines

Multi-drug resistance gene 1 (*MDR1*) encodes the P-glycoprotein protein that plays a role in the efflux of chemotherapeutic agents such as taxol, tamoxifen, and doxorubicin. To determine the potential of dichapetalin M to downregulate expression of *MDR1*, RT-qPCR was performed as described above, by comparing *MDR1* expression in dichapetalin M-treated cells versus the control vehicle treatment.

### Statistical Analysis

Statistical analysis was carried out using GraphPad Prism (version 7). Based on the number of groups, one-way ANOVA or a t-test was performed to evaluate statistical significance.

## Results

### Selectivity of Dic M for ER+ breast cancer cell lines and its ER agonistic effects

The selectivity of dichapetalin M for breast cancer cell lines with different receptor statuses was investigated using an MTT assay following 48 and 72-hour treatment. The IC_50_ values obtained after treatment were compared to assess selectivity across the cell lines. Dic M exhibited higher cytotoxicity towards MCF-7 than MDA-MB 231 and MDA-MB 453, both of which are ER-negative (**Table 2)** (Dai *et al*., 2017). Results obtained indicated that dichapetalin M is selective for the ER+ breast cancer cell line compared with cells lacking ER expression or expressing other receptors.

**Table 2:** The IC_50_ of Dichapetalin M in MCF-7, MDA-MB 231 and MDA-MB 231 after 48hrs and 72hrs of treatment.

| Cell lines | $IC_{50}$ of Dichapetalin M values at different time points (μM) | |
| --- | --- | --- |
|  | 48hrs | 72hrs |
| MCF-7 | 4.71±1.20 | 3.95 ± 1.43 |
| MDA-MB 453 | 11.07± 5.13 | 8.73 ± 5.34 |
| MDA-MB 231 | 23.84 | 36.77 |
| Selectivity to ER- Her 2<sup>+</sup>/ ER<sup>+</sup> | 2.35 | 2.21 |
| Selectivity to ER- / ER | 5.06 | 9.30 |

To investigate the agonistic effect of Dic M, the expression of *AREG* (amphiregulin), a critical effector molecule of ER signaling was monitored after treating MCF-7 (an ER-positive breast cancer cell line) with Dic M. Treatment of cells with gradient concentrations of Dic M, Estradiol (ER agonist) and co-treatment with E2 and 25% Dic M showed a dose-dependent increase in the expression of *AREG*. The expression of *AREG* increased by 2-fold and 3-fold following treatment with 25% dichapetalin M and E2, respectively. Co-treatment of cells with both E2 and 25% dichapetalin M (0.99 μM) showed a strong agonistic effect with a fold induction of about 12 after 24 hours of treatment (**Figure 2**).

**Figure 2:**
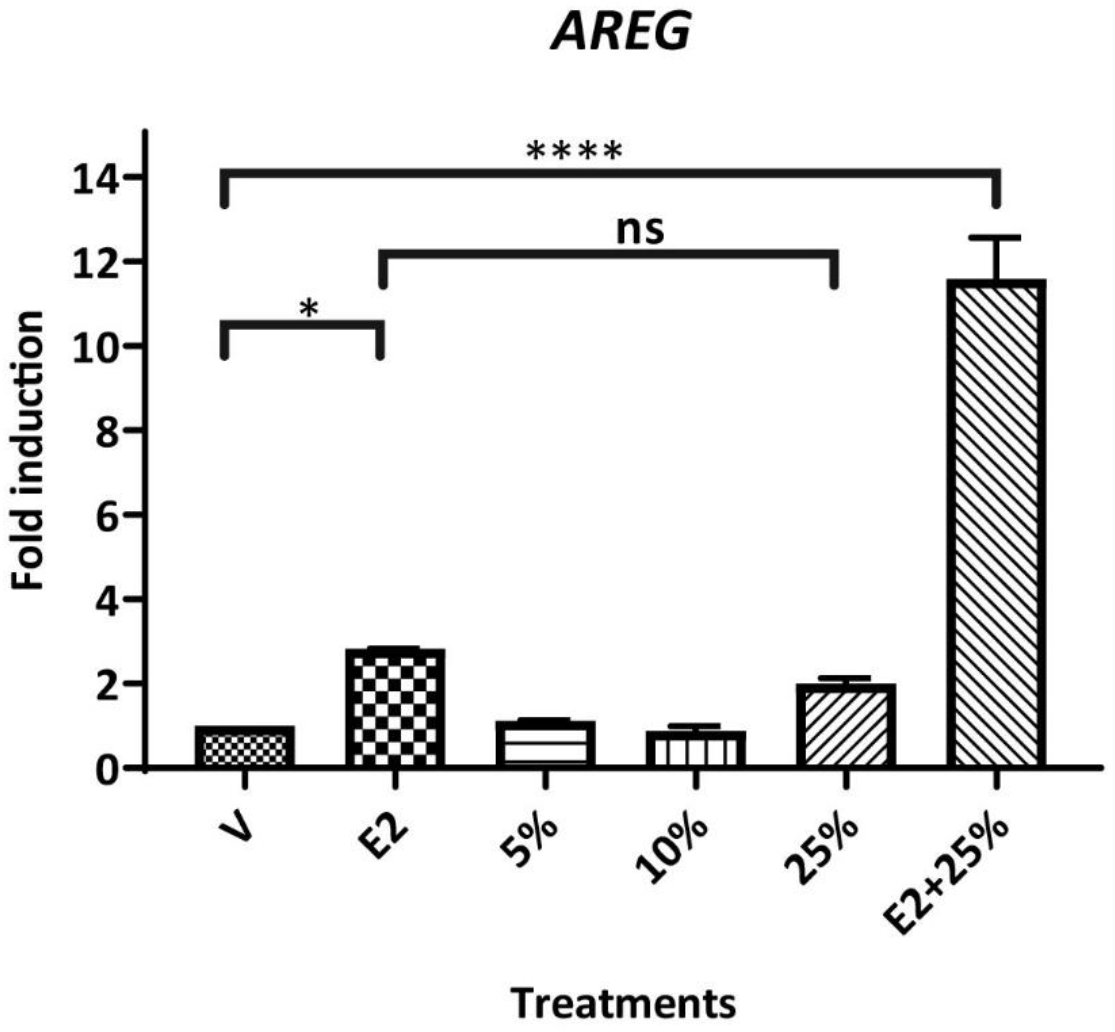
Effect of Dic M on the expression of Amphiregulin (*AREG*) in MCF-7. Cells were exposed to Dic M (at 5%, 10%, and 25% of its 72 hrs IC50) for 24 hrs. Cells treated with 0.01% DMSO, 10nM estradiol (E2) were used as vehicle only (V) and the ER agonist, respectively. Cells were co-treated with E2 and 25% Dic M to assess the antagonistic or synergistic effect of Dic M. ****, p< 0.0001; ***, p = 0.0001 to 0.001; **, p = 0.001 to 0.01; *, p = 0.01 to 0.005; p-value > 0.05 was assigned ns-not significant.

### Effect of Dic M on actin polymerization

Treatment of cells with dichapetalin M after 24 hrs did not show any significant effect on the cell morphology or induce apoptosis (**Figure S1)**. However, actin integrity studies indicated that dichapetalin M disrupts actin polymerization in MCF-7 cell lines, leading to shortened and disorganized microfilaments in a concentration-dependent manner (**Figure 3A-B**). For instance, it was observed that cells treated with 25% Dic M inhibited actin polymerization in MCF-7 at both treatment times (24hrs and 48hrs) than cells treated with 10% Dic M, and there was no significant difference between the actin integrity of cells treated for 24 hrs and 48 hrs. It was also observed that treatment of cells with E2 increased actin polymerization compared to cells treated with Dic M as can be seen in **Figure 3C**. However, Dic M could not significantly inhibit actin polymerization in the presence of ER agonist estradiol (E2), a ligand implicated in actin polymerization (**Figure 3C, panels 2 and 3)** after 24 hrs of treatment.

**Figure 3:**
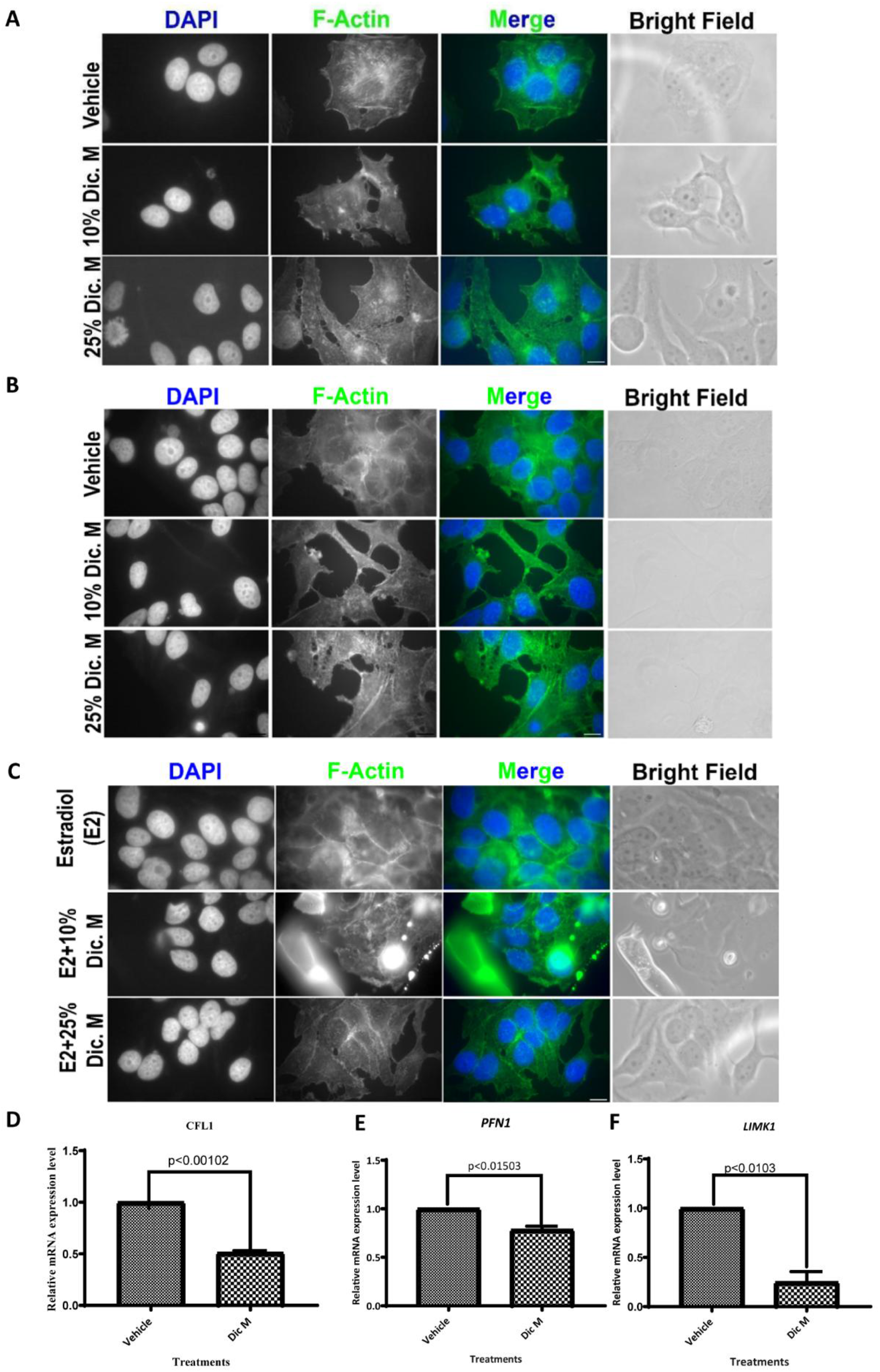
Effect of Dic M on Actin polymerization. A-B) Representative images showing actin integrity in MCF-7 after various treatments at 24 hrs and 48hrs. C) Representative images showing actin integrity in MCF-7 after co-treatment with E2 and Dic M. Images were taken at x100 (Pixel=30nm). Nucleus stained with DAPI (in blue) and F-actin stained with Phalloidin conjugated with a green fluorescent Alexa Fluor-488 dye (in green). D) mRNA expression levels of CFL1, PFN1 and LIMK1 after treatment with 25% Dic M; 0.01% DMSO was used as vehicle control. Samples without cDNA template were used as negative controls, and β-ACTIN was used as the reference gene. The data was analyzed statistically using t-test and setting the p-value at < 0.05.

Results from the actin polymerization studies were validated by RT-qPCR to observe the effect(s) of dichapetalin M on the expression of *CFL1, PFN1*, and *LIMK1*, genes that promote actin polymerization. The expression studies were performed after 24 hrs with 25% Dic M treatment. Cells treated with 0.01% DMSO only (vehicle) served as the control. Results indicate that Dic M down-regulates *CFL1, PFN1* and *LIMK1* expression significantly.

### Effect of Dic M on cell migration

A scratch-wound healing assay was performed to assess the effect of DIc M on cell migratory potential. The results of the scratch-wound healing assay demonstrate that after 24 hrs of treatment, there was complete wound closure. However, the rate of wound closure (area/time) was higher in cells treated with vehicle (0.01% DMSO) at different time points compared with cells treated with varying concentrations of Dic M. The rate of wound closure inhibition was not observed to occur in a dose-dependent manner (**Figure 4A-B**).

**Figure 4:**
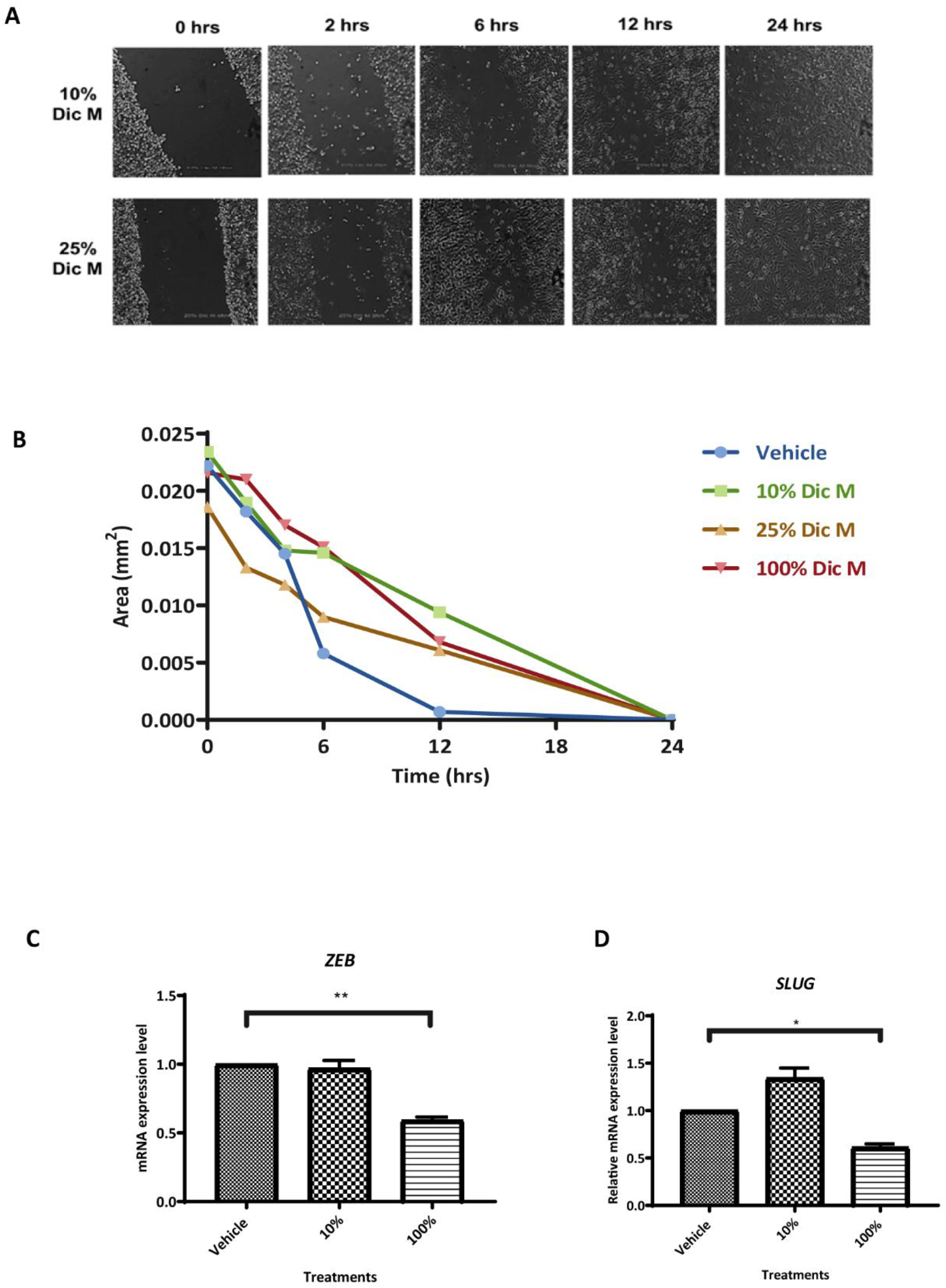
Effect of Dic M on MDA-MB 231 migration. A) Representative images of the scratch wound assay. Images were taken at ×10 using an Optika light microscope. B) Shows changes in the area of the wound at different time points after treatments, which is a measure of the rate of cell migration, and images were analyzed using Image J. C-D) The mRNA levels of ZEB and SLUG were determined using RT-qPCR. Data is presented as mean ± SEM of three independent experiments performed in triplicate. ** represents a p-value < 0.01, *represents a p-value < 0.05, and ns represents non-significant. The data was further analyzed statistically using a one-way ANOVA and Tukey’s multiple comparison test in GraphPad Prism (version 7).

The inhibition of cell migration was confirmed by observing the effect of Dic M on SLUG and ZEB, two repressors of E-cadherin: a protein whose down-regulation enhances epithelial-to-mesenchymal transition (EMT). Results obtained suggested that at higher concentrations of Dic M, *SLUG* and *ZEB*, repressors of E-cadherin were significantly down-regulated (**Figure 4C-D**).

### Effects of Dic M on the expression of apoptotic genes

Most cytotoxic compounds mediate their effects through the induction of apoptosis (Pérez-Soto *et al*., 2019). Therefore, to elucidate the mechanism by which dichapetalin M mediates its cytotoxic effect, its impact on the expression of pro-apoptotic and anti-apoptotic genes was examined. The results indicated that dichapetalin M significantly upregulated pro-apoptotic genes *BAD* (∼ 1.5-fold) and *BIM* (2-fold), however, *BCL-2* expression was unaffected by (**Figure 5**).

**Figure 5:**
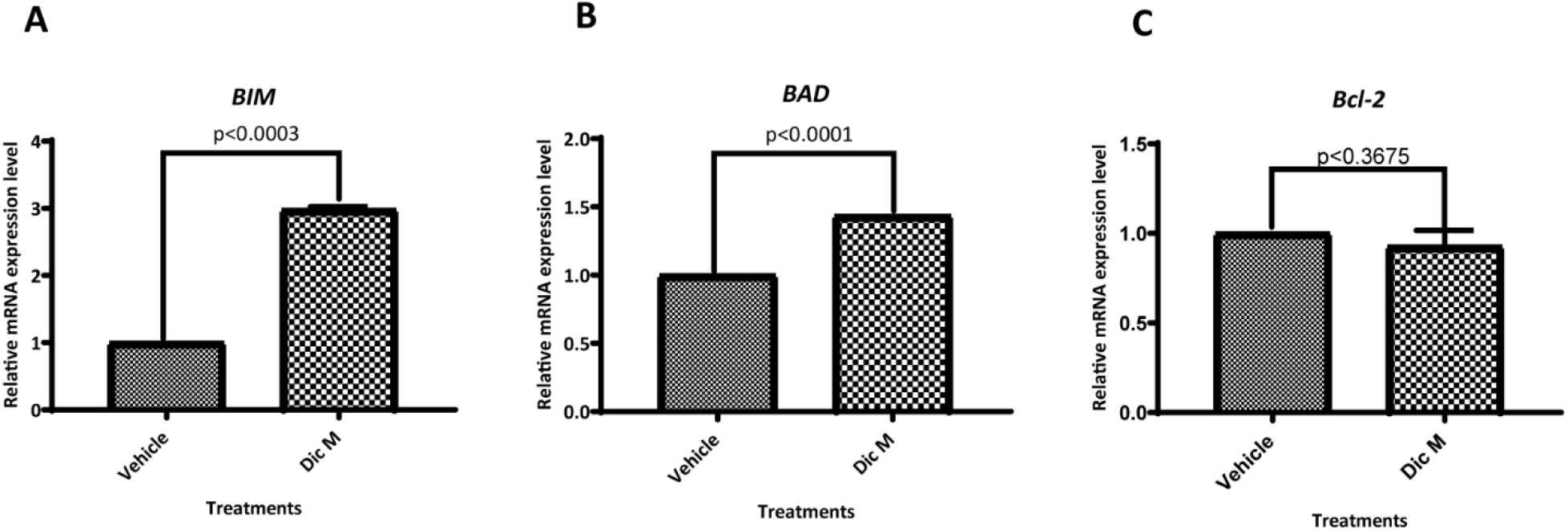
Effects of Dic M on the expression of apoptosis-related genes. A-C) mRNA expression level of BIM, BAD, and Bcl-2 after treatment with 25% dichapetalin M (Dic M); 0.01% DMSO was used as a vehicle. Samples without cDNA template were used as negative controls, and β-ACTIN was used as the reference gene. The data was analyzed statistically using t-test and setting the p-value at < 0.05.

**Figure 6:**
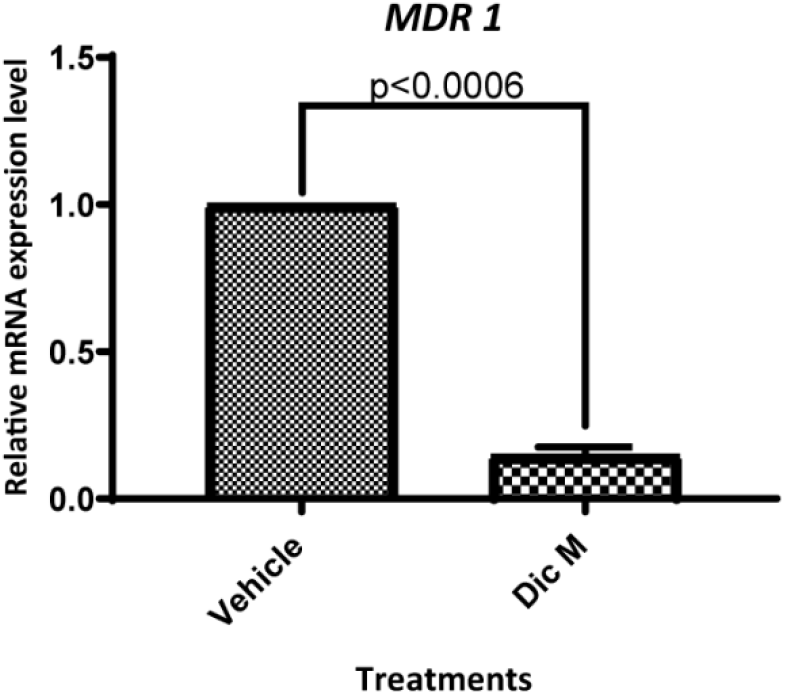
Effect of Dic M on MDR1 expression in the MCF-7 cell line. Gene expression studies on MDR1 after treatment of MCF-7 cells with 25% Dic M 0.01% DMSO was used as a vehicle control. Samples without cDNA template served as negative controls and β-actin was used as the reference gene. The data was analyzed statistically using t-test and setting the p-value at < 0.05.

**Figure 7:**
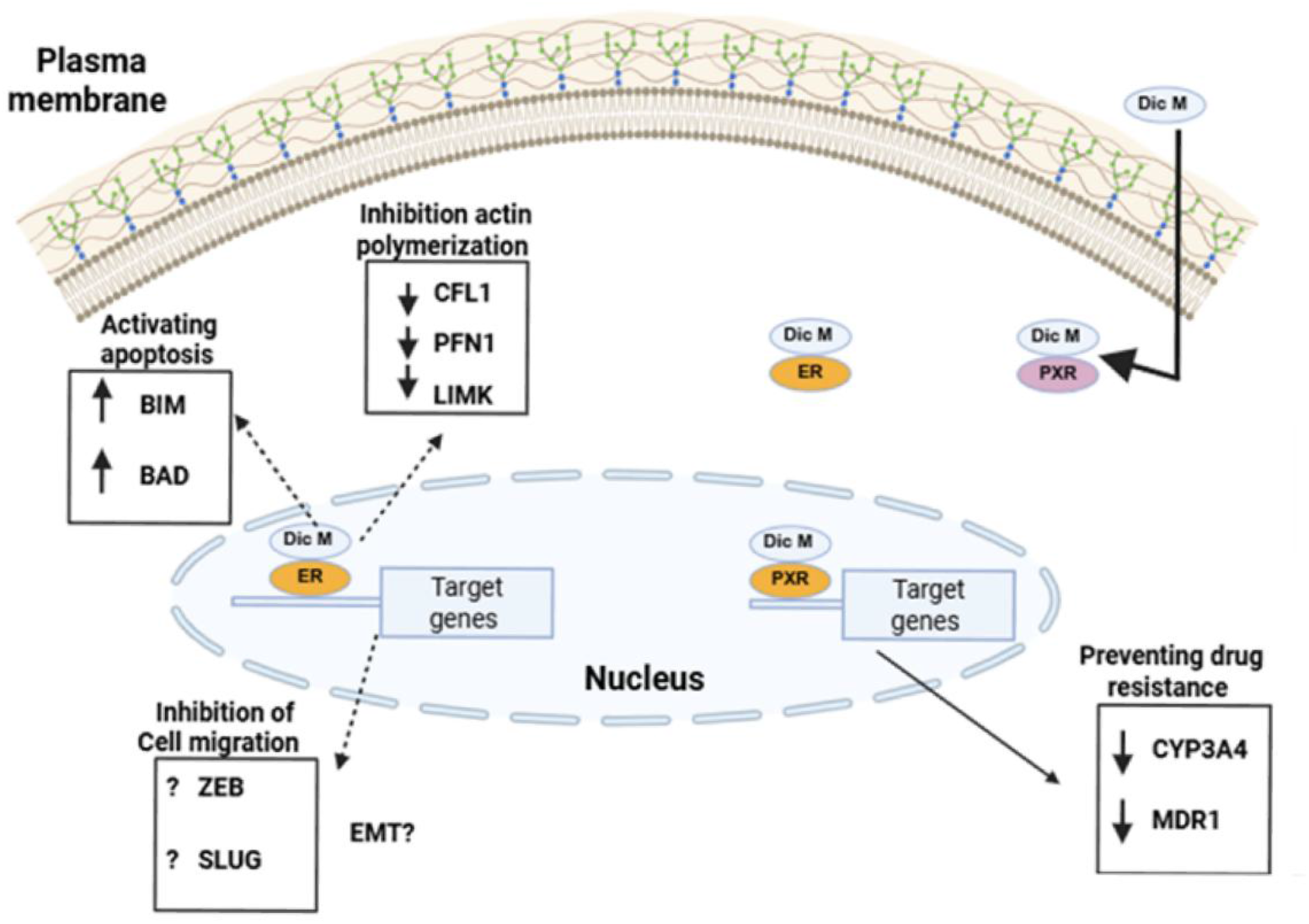
Proposed mechanism of action for Dic M. Dichapetalin M (Dic M) diffuses through the plasma membrane due to its hydrophobic moiety and interacts with the ER receptor. This interaction through an unknown pathway result in the upregulation of proapoptotic genes such as BAD and BIM, possibly accounting for the cytotoxic effect of the compound. Dic M treatment downregulated *LIMK1, CFL1* and *PNFL1*, reported to enhance normal actin polymerization, possibly resulting in the F-actin disorganization observed. Although Dic M was observed to slow down the rate of cancer cell line migration, it downregulated the expression of EMT-related genes, ZEB and SLUG at high doses. Dic M downregulated the expression of MDR1, a gene regulated by PXR and implicated in breast cancer resistance.

### Effects of Dic M on the expression of multidrug resistance gene 1 *MDR1*

To examine the ability of Dic M to prevent drug resistance in cancer cells, the compound’s effect on the expression of multidrug resistance gene 1 (*MDR1)* was investigated. *MDR1* encodes a P-glycoprotein that mediates the efflux of most therapeutic agents used in cancer treatment. Dichapetalin M downregulated the expression of *MDR1* compared to that of the vehicle-only treatment.

## Discussion

Chemotherapy remains the most preferred method for the treatment of breast cancer. While chemotherapy enhances tumor control and increases survival rates, its associated side effects, like drug resistance and non-specificity, can negatively impact patient adherence to treatment, potentially leading to poorer outcomes. For instance, the prolonged usage of tamoxifen, which is currently used for the treatment of ER+ breast cancer, fails to address the issues of breast cancer resistance and side effects such as impaired bone development in patients (Zidan et al., 2004; Chang et al., 2011). In our study, dichapetalin M, previously reported to be cytotoxic against certain carcinoma cell lines, was tested for its antiproliferative potential in breast cancer cell lines. *In silico* studies previously identified that dichapetalin M binds to the estrogen receptor (ER) and the Pregnane-X-receptor (PXR). The interaction of dichapetalin M with PXR was previously verified by our work in Chama *et al*. (2021). However, its interaction with ER was yet to be verified.

In this study, we elucidated the mechanism of action of Dic M by first verifying its interaction with ER. This was achieved by assessing the selectivity of Dic M in inducing its cytotoxic effect in an ER-positive cell line MCF-7, relative to ER-negative cell lines MDA-MB 231 and MDA-MB 453. The selectivity index obtained by comparing the IC_50_ of Dic M in MCF-7 with that in the ER-negative cell lines was >2 (Table 2), suggesting high selectivity of Dic M to the estrogen receptor. Additionally, Dic M in this study acted as an agonist to ER in a dose-dependent manner, such that cells treated with higher doses showed a more robust increase in the expression of *AREG* (amphiregulin) (Figure 2). *AREG* expression increased exponentially upon co-treatment of MCF-7 with DIc M and E2, suggesting that E2 and Dic M have a synergistic effect on ER signaling. This result is in contrast with the effect of a known ER antagonist, fulvestrant that bears a similar structure to Dic M (Osborne et al., 2004).

Given that ER signaling has been linked to actin polymerization (Lachowski et al., 2020)-a process known to contribute to cancer cell migration-the impact of Dic M on actin polymerization in MCF-7 cells was further investigated. Dic M inhibited actin polymerization in MCF-7 in a dose-dependent manner. Results were, however, not time dependent. Co-treatment of MCF-7 cell lines with dichapetalin M and E2 attenuated Dic M-induced actin disorganisation. Based on the effect of Dic M on the expression of factors involved in regulating actin turnover, Dic M potentially disorganises actin by downregulating the expression of the actin-polymerization-promoting factors *CFN1*, resulting in a commensurate decline in the expression of *LIMK1* that requires *CFN1* to execute its inhibitory effect on actin polymerization. These results correlated with previous work which showed a correlational decrease in the expression of *CFL1* during F-actin disorganization in PC-3, DU-145 and MDA-MB-231 cells following bisphosphonate alendronate treatment, a bisphosphonate derivative, for the treatment of bone diseases and bone metastasis (Virtanen *et al*. 2018). The expression of *PFN1*, which helps recycle G-actin monomers to ensure actin polymerization, was also down-regulated.

Breast cancer cells can migrate from their primary site to a distant organ with a suitable microenvironment that supports their growth and development. As part of elucidating its mechanism of action, we investigated the effect of Dic M on cancer cell migration. The results obtained from the scratch-wound assay indicated that Dic M slows down the rate of cell migration in MDA-MB 231, similar to a naproxen derivative, which was reported to impair cell migration in breast cancer cell lines (Deb et al., 2014). The inhibition of cell migration was assessed by observing the effect of Dic M on *SLUG* and *ZEB*, two repressors of E-cadherin: a protein whose down-regulation enhances epithelial-to-mesenchymal transition (EMT) (Adhikary et al.,2014). *SLUG* and *ZEB* expression were significantly downregulated upon treatment with higher doses of Dic M (100% of IC_50_ of dichapetalin M after 72 hrs of treatment). This result is no different from research conducted by Hsieh *et al*. (2019), which linked the decrease in *CFL1* expression with inhibition of cell migration in breast cancer cell lines upon treatment with the anticancer drug paclitaxel (Hsieh et al.2019).

Dichapetalin M also mediated its cytotoxic effect via the apoptotic pathway by significantly (*p*<0.05) up-regulating the pro-apoptotic genes *BIM* and *BAD*. Dic M did not affect the expression of the anti-apoptotic gene *BCL-2* (Figure 5). Work done by Leung & Wang, (2000) in assessing the effect of genistein, a phytoestrogen, on *BCL-2* expression during apoptosis in MCF-7, showed that *BCL-2* expression remained similar to the control even at concentrations that induced maximum cell death (Leung & Wang, 2000). This suggests that Dic M may exert its cytotoxic effect through a pathway similar to that of known phytoestrogens.

To explore dichapetalin M as a potential drug candidate for the treatment of resistant breast cancer, its effect on the expression of *MDR1*, which encodes a key protein involved in the efflux of several drugs, including some anti-breast cancer drugs, was examined. Our results indicated that dichapetalin M significantly downregulates the expression of *MDR1* in MCF-7. These findings further support the antagonistic activity of dichapetalin M against PXR, as previously reported by us (Chama *et al*. 2021). The results are also consistent with earlier studies showing that SPA70, a known PXR antagonist, downregulated the expression of both CYP3A4 and MDR1 proteins (Lin et al., 2017). Demonstrating that in contrast to tamoxifen, which upregulates *MDR1* expression and contributes to resistance in breast cancer therapy, Dic M downregulates its expression (Nagaoka et al., 2006).

## Conclusions

Using an *in vitro* target validation approach, we have demonstrated that Dic M acts as a selective ER agonist and putatively interacts with ER. The cytotoxic effect of Dic M was mediated via the upregulation of pro-apoptotic genes and downregulation of an anti-apoptotic gene. Its anti-migrastatic potential was due to its ability to disorganize actin-polymerization through the downregulation of *CFL1*, which enhances cell migration. DIc M, therefore, has the potential to be used for the treatment of ER+ breast cancer with reduced resistance via downregulation of *MDR1*. The evidence provided in this study gives some leads on the targets and mechanism of action of Dic M. However, investigating the effect of Dic M on global gene expression would further elucidate the mechanisms of action of the compound. Selectivity assays that will determine its suitability for breast cancer cells would be crucial. Finally, *in vivo* studies would determine the toxicity of dichapetalin M and further solidify its potential application in the treatment of ER+ resistant breast cancer.

## Abbreviations

Dic M: dichapetalin M
ER: Estrogen receptor
PXR: Pregnane X receptor
MDR: Multidrug-resistant
ERα: Estrogen receptor alpha subtype
ERβ: Estrogen receptor beta subtype
EGFR: Epidermal growth factor receptor
DMEs: Drug Metabolism enzymes
PR: Progesterone receptor
EMT: Epithelial-to-mesenchymal transition
4-OH tamoxifen: 4-hydroxy tamoxifen.

## Data Availability

All data generated or analyzed during this study are included in the manuscript. For further inquiries contact the corresponding author.

## Authors contribution

G.Y., L.P. and M.A.C. conceived the project. G.Y. performed experiments. G.Y. and L.P. analyzed data.

K.Y.A. validated the findings. L.P. and M.A.C. provided the reagents. G.Y. and L.P. wrote the paper. G.Y., K.Y.A., M.A.C. and L.P. edited and reviewed the paper. L.P. supervised the project.

## Acknowledgement

We want to acknowledge Jessica Asomaniwaa Armah who isolated dichapetalin M under the supervision of Prof. Chama. We are grateful to members of Paemka Genetic/Genomic laboratory (both past and present), Chama lab and Appiah-Opong lab, especially Isaac Tuffour at Noguchi for their support and expertise that greatly enhanced this work. Finally, we are grateful to CAPREx/Alborada research Grant [Grant Number RG86330] and Isaac Newton Matching Fund [Grant Number 17.07(c)] which supported the isolation of dichapetalin M. Finally, to WACCBIP for the award of $8300 which included a research grant of $5000.

## Supplemental data

**Figure S1:**
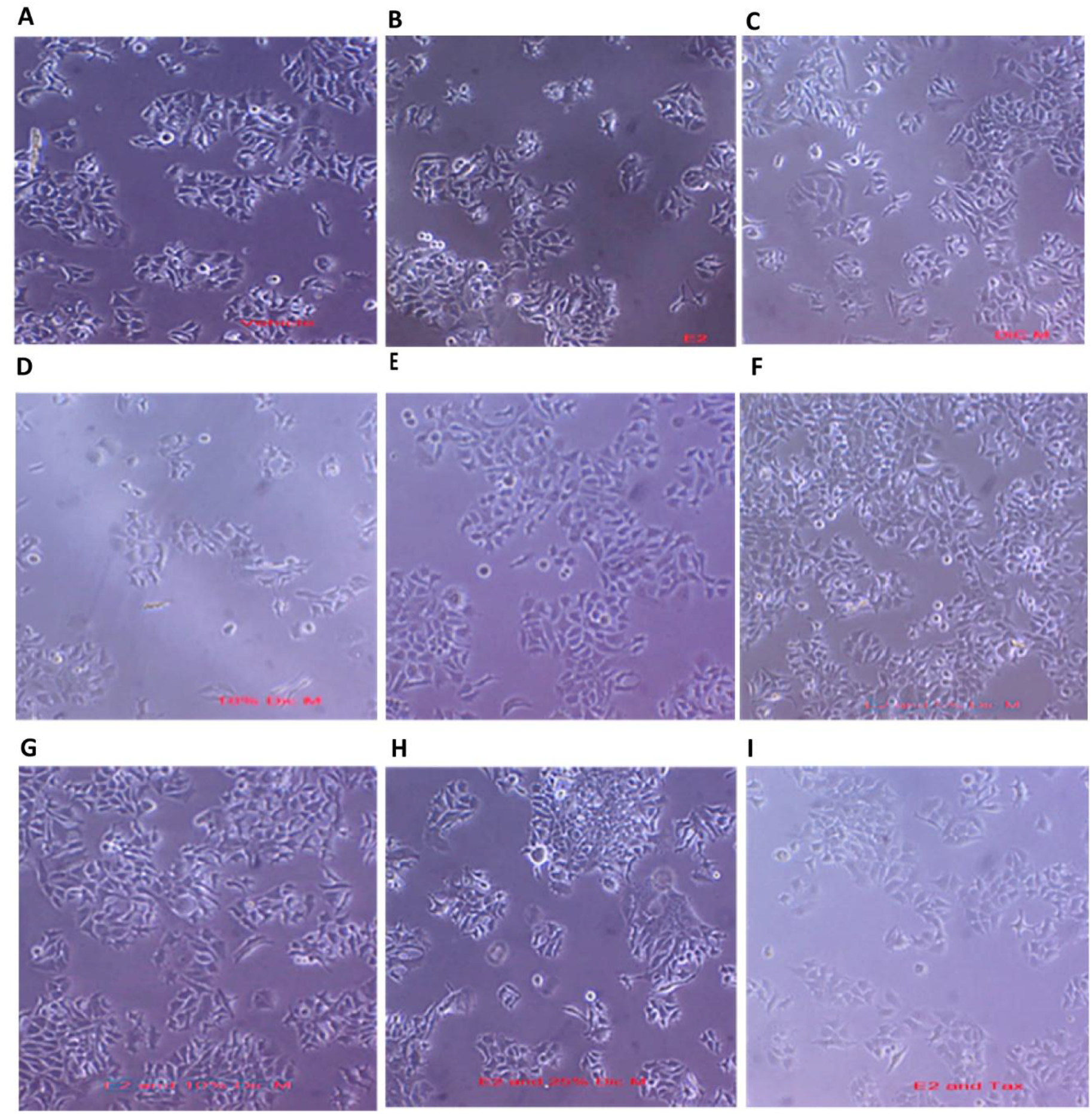
Phase-contrast photomicrographs of cultured MCF-7 cells with various treatments for 24 hours. Images obtained at x10 after treating of 1.0 × 10^5^ cells/ml cultured on a 22mm × 2mm with 0.01% DMSO vehicle (A), E2 (B), 5%, 10% and 25% Dichapetalin M (C, D and E respectively), and co-treatment with E2 and (5%, 10%, 25% Dichapetalin M and 10µM tamoxifen) (F, G, H and I, respectively). Images show healthy cells after 24 hrs of treatment with compounds

**Figure S2:**
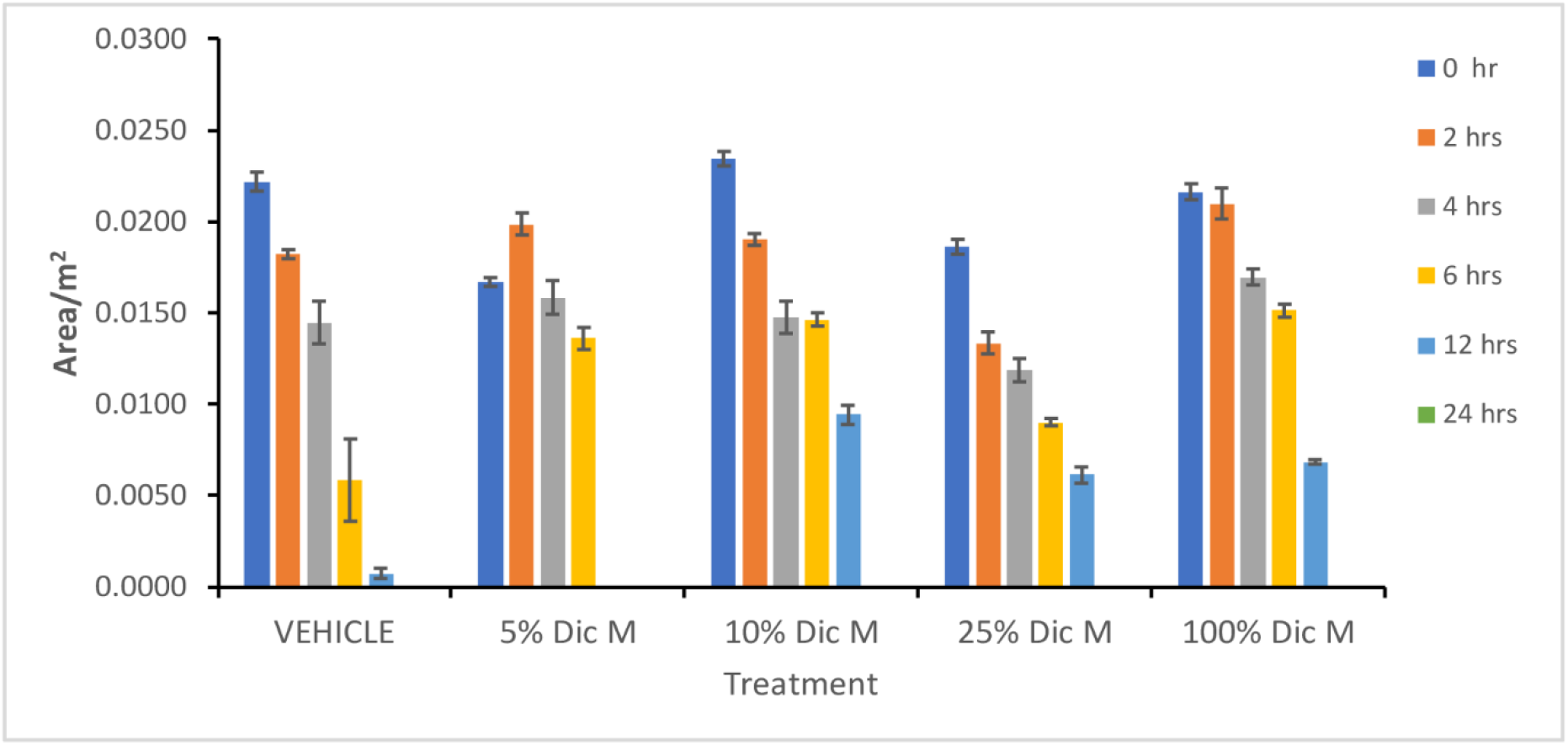
Bar chart showing effect of varying concentrations of Dichapetalin M on cell migration by measuring the scratch-wound area over time.

**Table S1:** Concentration of Dichapetalin M used for mechanistic studies.

| % of $IC_{50}$ | Actual concentration/μM |
| --- | --- |
| 5% | 0.20 |
| 10% | 0.40 |
| 25% | 0.99 |
| 100% | 3.95 |

**Table S2:**
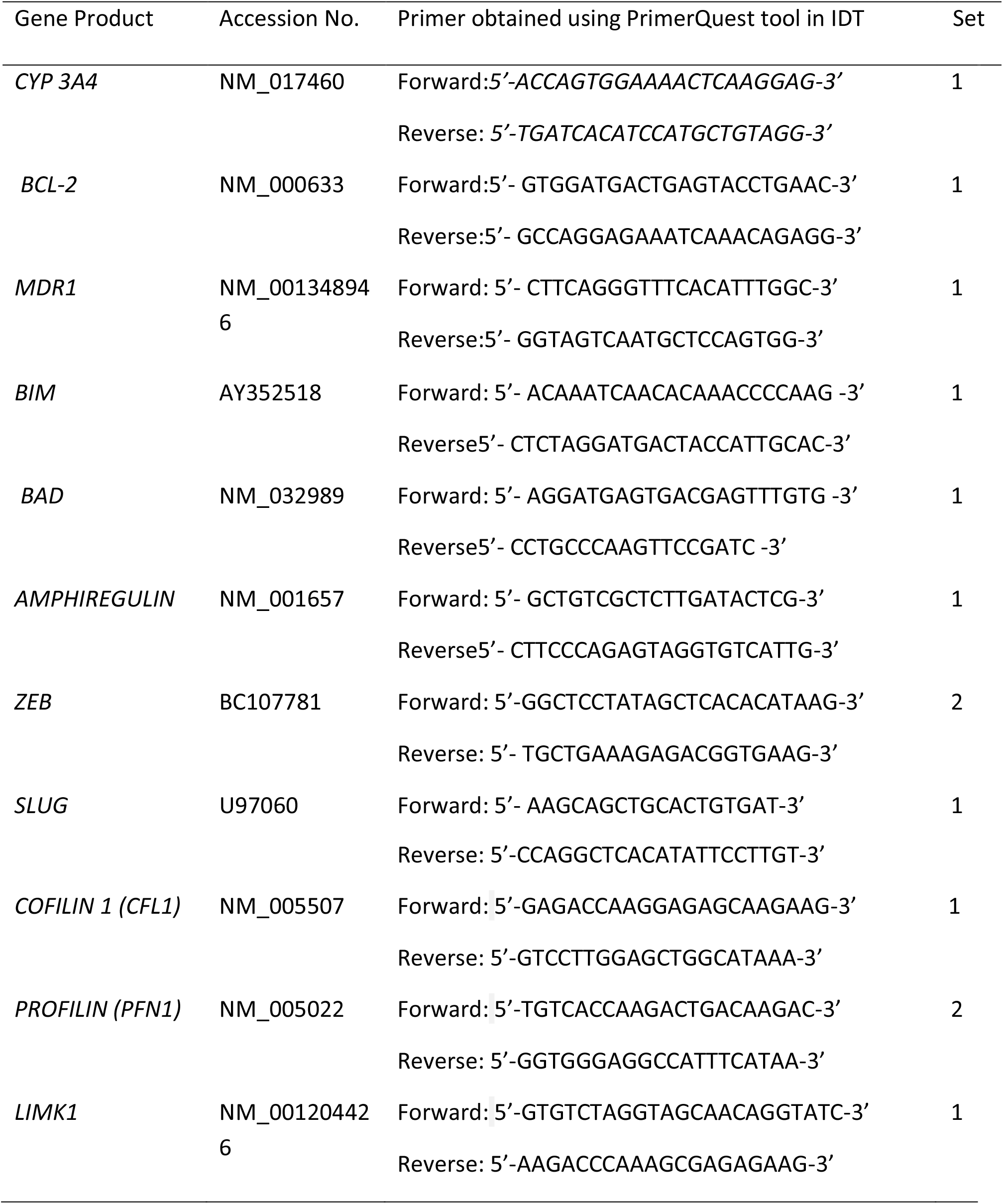
Primers for qRT-PCR.

## References

Abachi, M., Salati, M., Araghi, S., Shirkoohi, R., & Eslamifar, A. (2017). Molecular analysis of Acquired Tamoxifen resistance in breast cancer cell line. Asian Pac. J. Cancer Biology, 2(2), 41–49

Adhikary, A., Chakraborty, S., Mazumdar, M., Ghosh, S., Mukherjee, S., Manna, A., Mohanty, S., Nakka, K. K., Joshi, S., De, A., Chattopadhyay, S., Sa, G., & Das, T. (2014). Inhibition of epithelial to mesenchymal transition by e-cadherin up-regulation via repression of slug transcription and inhibition of e-cadherin degradation dual role of scaffold/matrix attachment region-binding protein 1 (smar1) in breast cancer cells. J Biol Chem. 289(37), 25431–25444. 10.1074/jbc.M113.527267

Badisa, R. B., Darling-Reed, S. F., Joseph, P., Cooperwood, J. S., Latinwo, L. M., & Goodman, C. B. (2009). Selective cytotoxic activities of two novel synthetic drugs on human breast carcinoma MCF-7 cells. Anticancer res., 29(8), 2993–2996.

Bray, F., Laversanne, M., Sung, H., Ferlay, J., Siegel, R. L., Soerjomataram, I., & Jemal, A. (2024). Global cancer statistics 2022: GLOBOCAN estimates of incidence and mortality worldwide for 36 cancers in 185 countries. CA Cancer J. for Clin., 74(3), 229–263. 10.3322/caac.21834

Chama, M. A., Modos, D., Mervin, L. H., Owusu, K. B. A., Ayine-Tora, D. M., Egyir, B., Paemka, L., Yankson, G., Ohashi, M., Afzal, A. M., & Bender, A. (2021). Towards understanding antimicrobial activity, cytotoxicity and the mode of action of dichapetalins A and M using in silico and in vitro studies. Toxicon, 193, 28–37. 10.1016/j.toxicon.2021.01.002

Chang, B. Y., Kim, S. A., Malla, B., & Kim, S. Y. (2011). The effect of selective estrogen receptor modulators (SERMs) on the tamoxifen resistant breast cancer cells. Toxicol. Res., 27(2), 85–93

Chen, Y., Tang, Y., Wang, M. T., Zeng, S., & Nie, D. (2007). Human pregnane X receptor and resistance to chemotherapy in prostate cancer. Cancer Res., 67(21), 10361–10367. 10.1158/0008-5472.CAN-06-4758

Clusan, L., Ferrière, F., Flouriot, G., & Pakdel, F. (2023). A Basic Review on Estrogen Receptor Signaling Pathways in Breast Cancer. Int. J. Mol. Sci. (Vol. 24, Issue 7).

Dai, X., Cheng, H., Bai, Z., & Li, J. (2017). Breast cancer cell line classification and Its relevance with breast tumor subtyping. J. Cancer, 8 (16), 3131–3141. 10.7150/jca.18457

Deb, J., Majumder, J., Bhattacharyya, S., & Jana, S. S. (2014). A novel naproxen derivative capable of displaying anti-cancer and anti-migratory properties against human breast cancer cells. BMC Cancer, 14(1). 10.1186/1471-2407-14-567

Fang, L., Ito, A., Chai, H.-B., Mi, Q., Jones, W. P., Madulid, D. R., Oliveros, M. B., Gao, Q., Orjala, J., Farnsworth, N. R., Soejarto, D. D., Cordell, G. A., Swanson, S. M., Pezzuto, J. M., & Kinghorn, A. D. (2006). Cytotoxic Constituents from the Stem Bark of Dichapetalum gelonioides Collected in the Philippines, J. Nat. Prod., 69(3), 332–337. 10.1021/np058083q

Kaminska, K., Akrap, N., Staaf, J., Alves, C. L., Ehinger, A., Ebbesson, A., Hedenfalk, I., Beumers, L., Veerla, S., Harbst, K., Ehmsen, S., Borgquist, S., Borg, Å., Pérez-Fidalgo, A., Ditzel, H. J., Bosch, A., & Honeth, G. (2021). Distinct mechanisms of resistance to fulvestrant treatment dictate level of ER independence and selective response to CDK inhibitors in metastatic breast cancer. Breast Cancer Res., 23(1). 10.1186/s13058-021-01402-1

Lachowski, D., Cortes, E., Matellan, C., Rice, A., Lee, D. A., Thorpe, S. D., & del Río Hernández, A. E. (2020). G Protein-Coupled Estrogen Receptor Regulates Actin Cytoskeleton Dynamics to Impair Cell Polarization. Front. Cel. Dev. Biol., 8. 10.3389/fcell.2020.592628

Lee, J., Alqudaihi, H. M., Kang, M. S., Kim, J., Lee, J. W., Ko, B. S., Son, B. H., Ahn, S. H., Lee, J. E., Han, S. W., Kim, Z., Hur, S. M., Lee, J. S., & Chung, I. Y. (2020). Effect of Tamoxifen on the Risk of Osteoporosis and Osteoporotic Fracture in Younger Breast Cancer Survivors: A Nationwide Study. Front Oncol., 10. 10.3389/fonc.2020.00366

Leung, L. K., & Wang, T. T. (2000). Nutrient-Gene Expression Bcl-2 Is Not Reduced in the Death of MCF-7 Cells at Low Genistein Concentration 1. J. Nutr. (Vol. 130).

Lin, W., Wang, Y. M., Chai, S. C., Lv, L., Zheng, J., Wu, J., Zhang, Q., Wang, Y. D., Griffin, P. R., & Chen, T. (2017). SPA70 is a potent antagonist of human pregnane X receptor. Nat. Commun., 8(1). 10.1038/s41467-017-00780-5

Long, C., Aussagues, Y., Molinier, N., Marcourt, L., Vendier, L., Samson, A., Poughon, V., Chalo, P. B., Ausseil, F., Sautel, F., Arimondo, P. B., & Massiot, G. (2013). Phytochemistry Dichapetalins from Dichapetalum species and their cytotoxic properties. Phytochemistry, 94, 184–191. 10.1016/j.phytochem.2013.03.023

Mouritzen, M. V., & Jenssen, H. (2018). Optimized scratch assay for in vitro testing of cell migration with an automated optical camera. Journal of visualized experiments: JoVE, (138), 57691.

Nagaoka, R., Iwasaki, T., Rokutanda, N., Takeshita, A., Koibuchi, Y., Horiguchi, J., Shimokawa, N., Iino, Y., Morishita, Y., & Koibuchi, N. (2006). Tamoxifen activates CYP3A4 and MDR1 genes through steroid and xenobiotic receptor in breast cancer cells. Endocrine, 30 (3), 261–268. 10.1007/s12020-006-0003-6

Nathan, M. R., & Schmid, P. (2017). A Review of Fulvestrant in Breast Cancer. Oncol. Ther., 5(1), 17–29. 10.1007/s40487-017-0046-2

Osborne, C. K., Wakeling, A., & Nicholson, R. I. (2004). Fulvestrant: An estrogen receptor antagonist with a novel mechanism of action. Br. J. Cancer, 90, S2–S6. 10.1038/sj.bjc.6601629

Osei-Safo, D., A. M., Addae-Mensah I., & Waibel, R. (2012). The Dichapetalins - Unique Cytotoxic Constituents of the Dichapetalaceae. In Phytochemicals as Nutraceuticals - Global Approaches to Their Role in Nutrition and Health. InTech. 10.5772/27224

Osei-Safo, D., Dziwornu, G. A., Appiah-Opong, R., Chama, M. A., Tuffour, I., Waibel, R., Amewu, R., Addae-Mensah, I., & Newman, D. J. (2017). Constituents of the Roots of Dichapetalum pallidum and Their Anti-Proliferative Activity. Molecules, 22(4). 10.3390/molecules22040532

Patel, H. K., & Bihani, T. (2018). Selective estrogen receptor modulators (SERMs) and selective estrogen receptor degraders (SERDs) in cancer treatment. In Pharmacol. Ther. (Vol. 186, pp. 1–24). Elsevier Inc. 10.1016/j.pharmthera.2017.12.012

Pérez-Soto, E., Estanislao-Gómez, C. C., Pérez-Ishiwara, D. G., Ramirez-Celis, C., & del Consuelo Gómez-García, M. (2019). Cytotoxic effect and mechanisms from some plant-derived compounds in breast cancer. Cytotoxicity-definition, identification, and cytotoxic compounds.

Schneider CA, Rasband WS, Eliceiri KW. NIH Image to ImageJ: 25 years of image analysis. Nat. Methods. 2012 Jul;9(7):671–5. doi: 10.1038/nmeth.2089. PMID: 22930834; PMCID: PMC5554542.

Shaath, H., Elango, R., & Alajez, N. M. (2021). Molecular classification of breast cancer utilizing long non-coding rna (Lncrna) transcriptomes identifies novel diagnostic lncrna panel for triple-negative breast cancer. Cancers, 13(21). 10.3390/cancers13215350

Skandalaki, A., Sarantis, P., & Theocharis, S. (2021). Pregnane x receptor (Pxr) polymorphisms and cancer treatment. Biomolecules (Vol. 11, Issue 8). MDPI AG. 10.3390/biom11081142

Virtanen, S. S., Ishizu, T., Sandholm, J. A., Löyttyniemi, E., Väänänen, H. K., Tuomela, J. M., & Härkönen, P. L. (2018). Alendronate-induced disruption of actin cytoskeleton and inhibition of migration/invasion are associated with profilin downregulation in PC-3 prostate cancer cells. Oncotarget, 9(66), 32593.

Zidan, J., Keidar, Z., Basher, W., & Israel, O. (2004). Effects of tamoxifen on bone mineral density and metabolism in postmenopausal women with early-stage breast cancer. Med. Oncol., 21(2), 117–121. 10.1385/MO:21:2:117

